# Auditory Brainstem Mechanisms Likely Compensate for Self-imposed Peripheral Inhibition

**DOI:** 10.1101/2022.11.26.518056

**Authors:** Abigayle Peterson, Vijayalakshmi Easwar, Lindsey Powell, Sriram Boothalingam

## Abstract

It is well known that the medial olivocochlear reflex (MOCR) in the brainstem, part of the efferent network, inhibits the cochlear active gain mechanism. The upstream neural influence of this peripheral inhibition is less understood. When the MOCR is activated, responses generated in the cochlea and cortex undergo putative attenuation, yet the amplitude of responses generated in the brainstem are perplexingly unaffected despite decreased input from the periphery. Based on known neural circuitry, we hypothesized that the inhibition of peripheral input is compensated for by equivalent positive feedback in the brainstem over time. We predicted that the inhibition can be captured at the brainstem with stimuli shorter (1.5 s) than previously employed long durations (4 min) where this inhibition is diminished due to compensation. Results from 18 normal hearing human listeners support our hypothesis in that when the MOCR is activated, there is a robust reduction of responses generated at the periphery, brainstem, and cortex for short stimuli and that brainstem inhibition diminishes for longer stimuli. Our methodology and findings have implications for auditory disorders such as tinnitus, evaluation of efferent function, and provides a novel non-invasive window into potential gain compensation mechanisms in the brainstem.

## Introduction

Efferent neural networks fine-tune and regulate afferent sensory inputs. One such network at the level of the brainstem, the medial olivocochlear reflex (MOCR), modulates activity of one of the most peripheral auditory structures, the cochlear outer hair cells (OHCs). The OHCs actively amplify basilar membrane motion for low-level sounds. When activated, the MOCR inhibits this amplification process, thus turning down the cochlear gain. This reduction in cochlear activity is thought to be useful for signal detection in noise (Winslow and Sachs, 1987; Warren & Liberman, 1989) and protection against loud sounds (Rajan, 1988; Liberman, 1990; Lauer and May, 2011). While the peripheral consequences of this inhibition are well-understood from studies using measures such as otoacoustic emissions (OAEs) and auditory nerve compound action potentials (CAPs), the upstream neural influence remains unknown, especially in humans. As such, the goal of this study was to determine the central consequences of peripheral MOCR inhibition. Our motivation stems from the need to uncover the ecological relevance of MOCR inhibition. This requires an improved understanding of its effect along the auditory pathway at different timescales.

It is well-established that stimulus-driven peripheral responses such as CAPs and OAEs undergo robust attenuation when the MOCR is activated (Galambos, 1956; Warren & Liberman, 1989; Siegel and Kim, 1982; review: Guinan, 2006). However, current evidence perplexingly exhibits a disparity of MOCR influence in the central systems based on the location of response generation along the auditory neuraxis. For instance, endogenous components of cortical responses (e.g., auditory steady state responses [ASSR] elicited at 40 Hz and thalamocortical loop resonance in the gamma band), undergo considerable attenuation in the presence of putative MOCR activation (Özdamar & Bohórquez, 2008; Ross et al., 2005, 2012; Maki et al., 2009). However, sensory-driven neural responses that originate in the brainstem (e.g., ASSR elicited at 80 Hz and auditory brainstem response [ABR] wave V), appear unaffected under same testing conditions (Özdamar & Bohórquez, 2008; Maki et al., 2009). This brainstem immunity to MOCR effects, typically elicited by contralateral noise, remains unexplained. Here, we seek to clarify our hypothesis that brainstem-dominant neural responses show immunity because local feedback networks in the brainstem compensate for the peripheral inhibition.

The rationale for our hypothesis is rooted in (1) previously identified circuits that are capable of such compensation (Fujino & Oertel, 2001; Hockley et al., 2021), and (2) gain compensation that occurs for more extreme peripheral input loss due to pathologies such as cochlear ablation, deafferentation, and conductive loss (Chambers et al., 2016; Parry et al., 2019; Sheppard et al., 2019). We predict that if local feedback networks do compensate for the MOCR-mediated peripheral inhibition, a latency between stimulus presentation, peripheral inhibition, and complete compensation would be apparent. Based on the time (~400 ms) it takes for the MOCR inhibition on OHCs to stabilize (Boothalingam et al., 2021; Backus and Guinan, 2006), we speculated that the time taken for complete compensation will fall between 0.5 and a few seconds. To test this prediction, we concurrently measured peripheral (cochlear) and neural (brainstem or cortical) responses in short (1.5 s) and long (4 mins) click-trains. Our results support our hypothesis in that, responses at the periphery (OAEs at 40 and 80 Hz), brainstem (80 Hz ASSRs) and the cortex (40 Hz ASSRs) demonstrate robust inhibition in shorter durations, however, for longer durations, only the inhibition of brainstem responses diminishes. That is, the inhibition appears to be compensated for at the brainstem between 1.5 s and 4 mins. This approach likely provides a window into brainstem feedback circuits that are involved in enhancing the peaks of complex signals (e.g., speech) and possibly maintaining the homeostasis in response to a, self-imposed, decrease in auditory input from the periphery (Brown, 2011; Fujino & Oertel, 2001; Hockley et al., 2021). In addition, our non-invasive, approach offers a solution to evaluating auditory efferent function in patients with sensorineural hearing loss and offers a new perspective on the ecological relevance of the MOCR.

## Materials and Methods

### Participants

Twenty young, clinically normal-hearing adults participated in the study. Clinically normal hearing was established by an unremarkable otoscopic examination, bilateral hearing thresholds ≤20 dB HL at octave frequencies from 0.25 to 8 kHz (SmartAuD, Intelligent Hearing Systems [IHS], FL, USA), normal middle ear function as measured by tympanometry (Titan, Interacoustics, Denmark), measurable (magnitude greater than 0 dB with at least 6 dB signal-to-noise ratio [SNR]) distortion product OAEs (0.5-6 kHz at 65/55 dB SPL, SmartDPOAE, IHS, FL), and self-report of no neurological disorders. Two participants were rejected from analysis due to excessive noise in OAEs, reducing the number of participants to 18 (mean age ± standard deviation [SD]) = 22.3±3 years; 1 male). Participants were either offered extra credit for their participation or compensated at the rate of $10/hour. The study procedures were approved by the University of Wisconsin-Madison Health Sciences Institutional Review Board. Written consent was obtained from all participants prior to data collection.

### Stimuli

All stimuli were digitally generated in MATLAB (v2017b; Mathworks, MA, USA) at a sampling rate of 96 kHz and a bit-depth of 24. The stimuli used to elicit OAEs and ASSRs were click trains with click rates of either 40 or 80 Hz presented at 65 dB peak-to-peak (pp) SPL. Whereas the 40 Hz clicks elicit a predominantly cortical response, the 80 Hz clicks elicit a predominantly brainstem response (Bidelman, 2018; Herdman et al., 2002; Kuwada et al., 2002; review: Dimitrijevic and Ross, 2008). Although henceforth we will refer to 40 Hz vs. 80 Hz responses as synonymous with cortical vs. brainstem sources for brevity given previously established dominant neural sources, it is acknowledged that both scalp-recorded responses reflect multiple neural generators. The clicks were bandlimited between 0.8 and 5 kHz to focus the stimulus energy on frequency regions where the MOCR function is most prominent, when measured using OAEs (Lilaonitkul and Guinan, 2012; Zhao and Dhar, 2012). Bandlimited clicks were generated in the frequency domain using a recursive exponential filter (Zweig and Shera, 1995; Charaziak et al., 2020) and inverse Fourier-transformed to the time domain. The duration of the click was ~ 108 μs. Clicks were presented in positive and negative polarities to reduce potential contamination with stimulus artifact when averaging for ASSRs. Broadband noise (0.001 to 10 kHz) was presented at 60 dB SPL in the contralateral ear to elicit the MOCR. Both the ipsilateral clicks and the contralateral noise are illustrated in Fig. 1. In-ear calibration was performed for clicks to ensure the peak-to-peak level of the click stimulus was 65 dB ppSPL in all participants. Broadband noise was calibrated in an ear simulator (Type 4157, Bruel & Kjaer, Denmark).

**Figure 1:**
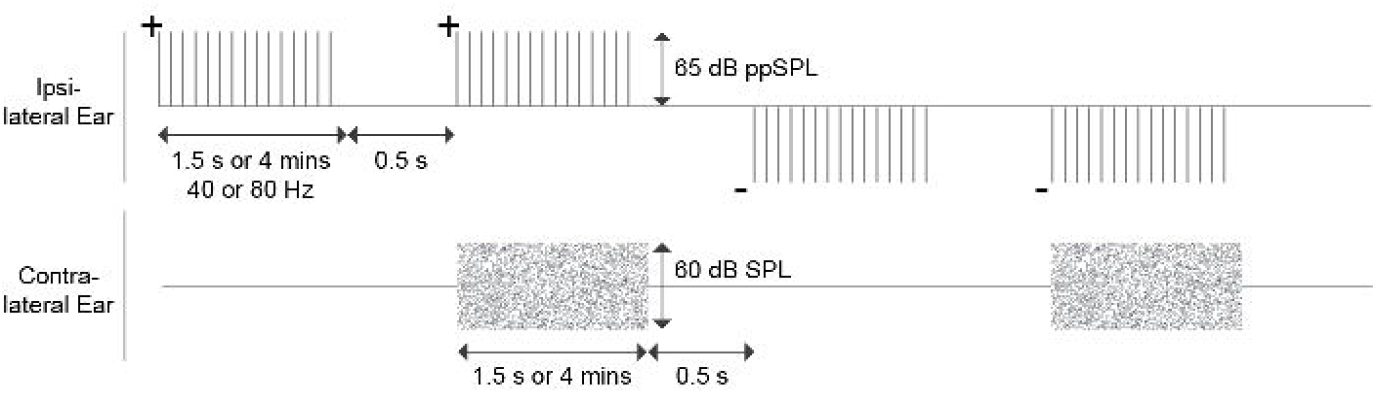
Schematic of the experimental protocol. Both 40 and 80 Hz clicks were presented in short (1.5 s) and long durations (4 mins) at 65 dB ppSPL, with and without a 60 dB SPL broadband noise in the contralateral ear. Clicks were presented in positive and negative polarities to facilitate ASSR averaging. Each stimulus duration was separated by 0.5 s of silence.

### Instrumentation

Stimuli were generated, delivered and controlled through an iMac computer (Apple, CA, USA) running Auditory Research Lab Audio Software (ARLas v4.2017; Goodman, 2017) on MATLAB. The iMac was interfaced with an external sound card (Fireface UFX+; RME, Germany) via thunderbolt for analog-to-digital-to-analog conversion at a sampling rate of 96 kHz. Clicks were presented in the ipsilateral ear via one of the miniature loudspeakers of the ER10C+ (Etymotic Research, IL, USA) system. Ear canal pressures were registered and amplified (+20 dB) by the ER10C+ probe microphone placed in each ear. To avoid changes in stimulus level over the course of the experiment, the probe placement was secured in the ear using foam tips and using silicone putty around the probe in-ear (Silicast, Westone Laboratories, CO; Boothalingam and Goodman, 2021). The MOCR eliciting broadband noise was presented in the contralateral ear using an ER2 (Etymotic Research, IL) insert earphone coupled with a foam tip of appropriate size.

Electroencephalography (EEG) amplitude was registered by the Universal Smart Box (USB; IHS, FL, USA) controlled by a PC equipped with the Continuous Acquisition Module (IHS, FL, USA) at the sampling rate of 10016 Hz. One of the IHS USB channels recorded triggers (5V impulses) that coincided with the onset of stimulus blocks to index EEG data accurately. A single-channel montage was used for EEG acquisition with three sintered Ag-AgCl electrodes. The vertex (Cz) was used as the non-inverting electrode site and the nape was used as the inverting site (Picton et al., 2003). The left collarbone was used as the ground. All electrode sites were cleaned with alcohol wipes and scrubbed with a mild abrasive gel (Nuprep) prior to affixing electrodes with adhesive sleeves and conduction gel (SignaGel). Electrode impedances were monitored throughout the experiment and were always <3 kΩ at each site.

### Experimental Design

The experiment was conducted in a double-walled sound-attenuating booth where participants sat comfortably in a recliner for the duration of the experiment. Participants were instructed to sit relaxed, not move, swallow as few times as comfortable during stimulation, and maintain a wakeful state (watching a silent, closed caption movie). As shown in Fig. 1, OAEs and ASSRs were measured at 40 and 80 Hz with and without contralateral noise, in short (1.5 s) and long (4 min) durations. The conditions (rate and duration) alternated between clicks with and without contralateral noise separated by 0.5 s of silence, and was repeated in positive and negative polarities. The long condition was a single block of 4 min recording completed separately for with- and without-contralateral noise, i.e., no repetitions. In contrast, in the short condition, 1.5 s-long click trains were repeated 160 times to match the total number of clicks presented in the respective long condition. This was done to avoid SNR differences in responses between long and short conditions analyzed in the frequency domain. The order of presentation in short/long conditions was counter-balanced but the 40 Hz short duration was always completed first to ensure maximum wakeful participant state as 40 Hz can be attenuated by sleep (Picton et al., 2003). The stimulus ear was chosen based on the ear with the largest DPOAE amplitude obtained during screening. The experimental procedure took about two hours with breaks, as desired by the participant, between conditions.

### Response Analysis

OAEs were extracted from click ‘epochs’, defined as the duration between two successive clicks. OAE analysis was performed offline in MATLAB using custom scripts. Raw ear canal pressure recording was bandpass filtered around the click frequency (0.8–5 kHz). An artifact rejection, where clicks with a root-mean-square (RMS) amplitude that fell outside the third quantile + 2.25 times the interquartile range (specific to the condition and within participants) were excluded from further analysis. Typically, less than 10% of the responses were rejected across participants. Data was grouped by duration with or without contralateral noise. The stimulus (0–4ms) and OAEs (6.5–12.5ms) were then separated for further analysis for both 40 and 80 Hz rates. Specifically, OAEs were used to determine the presence of the MOCR. The click stimulus was used to determine the presence of the middle ear muscle reflex (MEMR) that can potentially confound MOCR effects on OAEs (Boothalingam et al., 2018).

OAE amplitude was estimated as the RMS of ear canal pressure between 6.5–12.5 ms and the noise floor was estimated by the mean difference of two OAE RMS buffers (even and odd-numbered epochs). Prior to estimating the RMS amplitude, OAEs were considered in the frequency domain to extract OAEs 12 dB above the noise floor to ensure the MOCR-mediated inhibition was estimated only from high quality OAEs (Guinan, 2012; Goodman et al., 2012; Boothalingam et al., 2021). For each duration and rate condition, the RMS of OAE and stimulus amplitude (dB SPL) with noise were subtracted from the RMS without noise to compute the effect of MOCR and MEMR (in dB), respectively.

The same pre-processing strategy as ear canal pressure was applied to the raw EEG data. Raw EEGs were chunked into 1.5-s or 4-min epochs corresponding to short and long durations, respectively. Chunked EEGs were averaged over opposite stimulus polarities to minimize any stimulus artifacts. Finally, ASSR amplitudes were determined from the Fourier transforms of the averaged EEGs across with and without noise estimates at 40 and 80 Hz. EEG noise floor was estimated as the average of 8 frequency bins around the ASSR frequencies (Picton et al., 2003). The reduction in response amplitude with contralateral noise relative to no-noise condition is henceforth referred to as ‘inhibition’ in both OAEs and ASSRs.

### Middle ear muscle reflex (MEMR) estimation

When elicited by high level sound, the MEMR stiffens the ossicular chain, altering signal transfer through the middle ear (Boothalingam et al., 2021; Borg, 1968) and may thus confound MOCR effects on OAEs (Boothalingam et al., 2021; Goodman et al., 2013; Guinan et al., 2003). The click stimulus (0–4 ms) in the same frequency range as the OAEs was analyzed to determine the presence of MEMR. The same analyses as OAEs were applied to obtain stimulus level across conditions. To evaluate if the MEMR influenced OAEs, we performed two tests. First, a 3-way repeated measures analysis of variance (RM-ANOVA) was performed to test if the MEMR magnitude varied as a function of independent variables duration, click rate, and contralateral noise. Our results show that none of the main effects and interactions were significant (p>0.05). This suggests that the variables in the study did not systematically influence the stimulus level, therefore, even if any MEMR was activated, it did not influence our results. Second, Pearson correlations were performed to evaluate the relation between any change in stimulus amplitude on OAEs. As seen in Fig. 2, none of the correlations, except the long condition at 80 Hz, were significant suggesting that the changes in stimulus level did not influence changes in OAE level. For the significant correlation at long/80 Hz, removing the one outlier made the correlation non-significant (Fig. 2., panel C). Clearly, for this one participant, MEMR likely influenced the OAE change in this one condition. However, we did not exclude this data for further analysis below because (1) this is not consistent across conditions, (2) the influence is on the direction opposite to what is expected – increase in OAE level as opposed to a reduction, and (3) because group effects were not significant (3-way ANOVA). Taken together, the OAE changes presented in this study are driven by the MOCR and not the MEMR.

**Figure 2:**
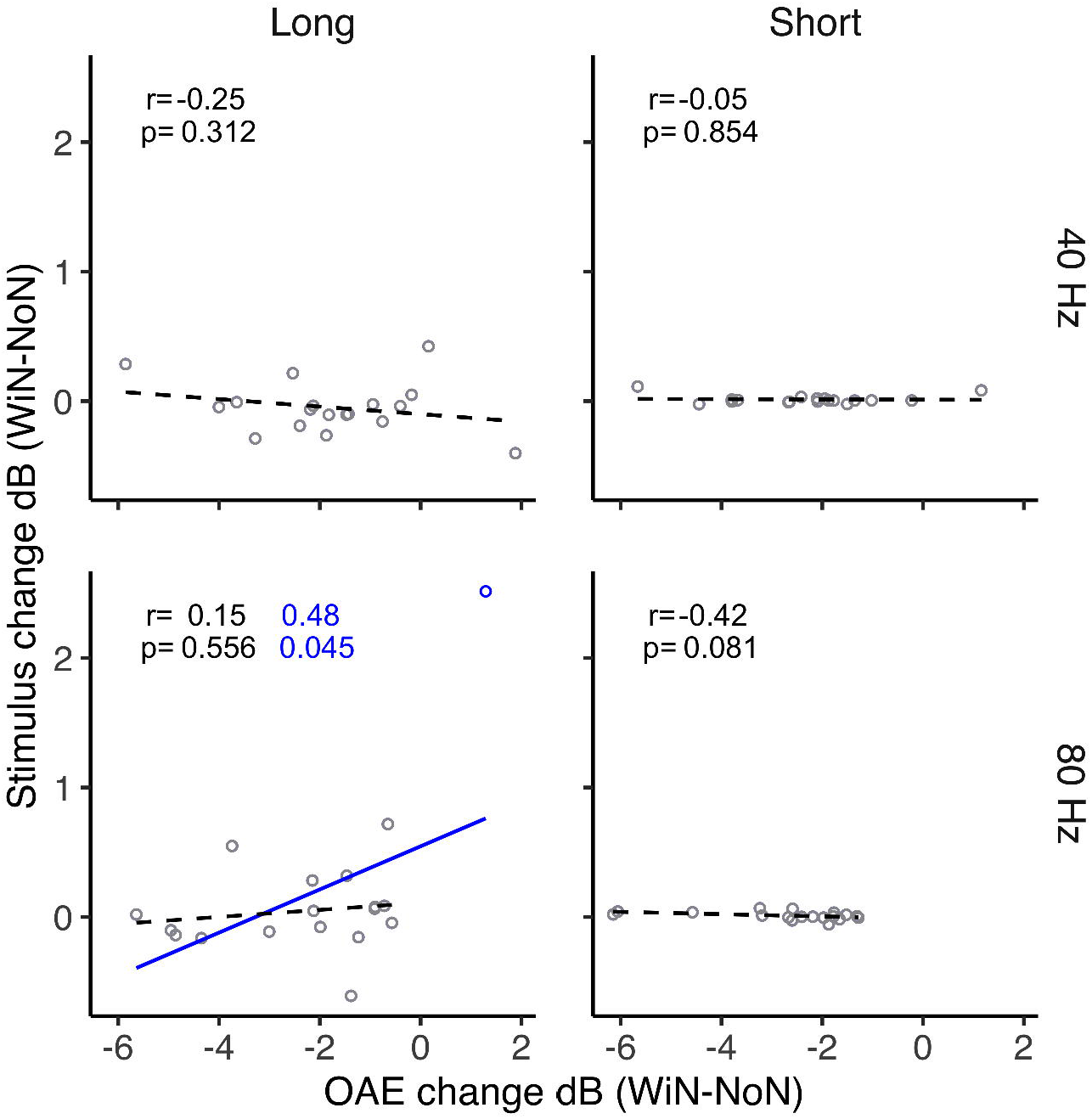
Stimulus vs. OAE change. Stimulus amplitude change as a function of OAE amplitude change for (A) 40 Hz click-rate, long stimulus duration (B) 40 Hz click-rate, short stimulus duration (C) 80 Hz click-rate, long stimulus duration (D) 80 Hz click-rate, short stimulus duration. Open circles represent individual participants. A black solid regression line represents a significant relationship between variables. A black dashed regression line represents a nonsignificant relationship between variables. The resulting correlation coefficient (*r*) and *p*-value are presented in each panel. In panel C, the blue solid regression line correlation when the outlier is included.

## Results

ASSR and OAE amplitudes are plotted as a function of click rate (40 vs. 80 Hz), contralateral noise (with vs. without), and duration (long vs. short) in Fig. 3. Notice the marked reduction in both OAE and ASSR amplitudes in the long condition. This reduction appears relatively unchanged for OAEs between short and long conditions. A 3-way repeated measures analysis of variance (ANOVA) for ASSRs revealed a significant three-way interaction. That is, the effect of noise varied as a function of duration and rate (*F*[1, 17])=33.28, *p*<0.001). Post-hoc *t*-tests were corrected for multiple comparisons using the False Discovery Rate (FDR; Benjamini and Hochberg, 1995). We report FDR corrected *p*-values and hence *p*<0.05 are to be interpreted as significant. These post-hoc tests demonstrate a significant effect of noise on 40 Hz ASSR in both short (*p*<0.001) and long (*p*=0.03) durations, however, the effect of noise on 80 Hz ASSR was only significant for the short (*p*<0.001) but not long (*p*=0.486) duration.

**Figure 3:**
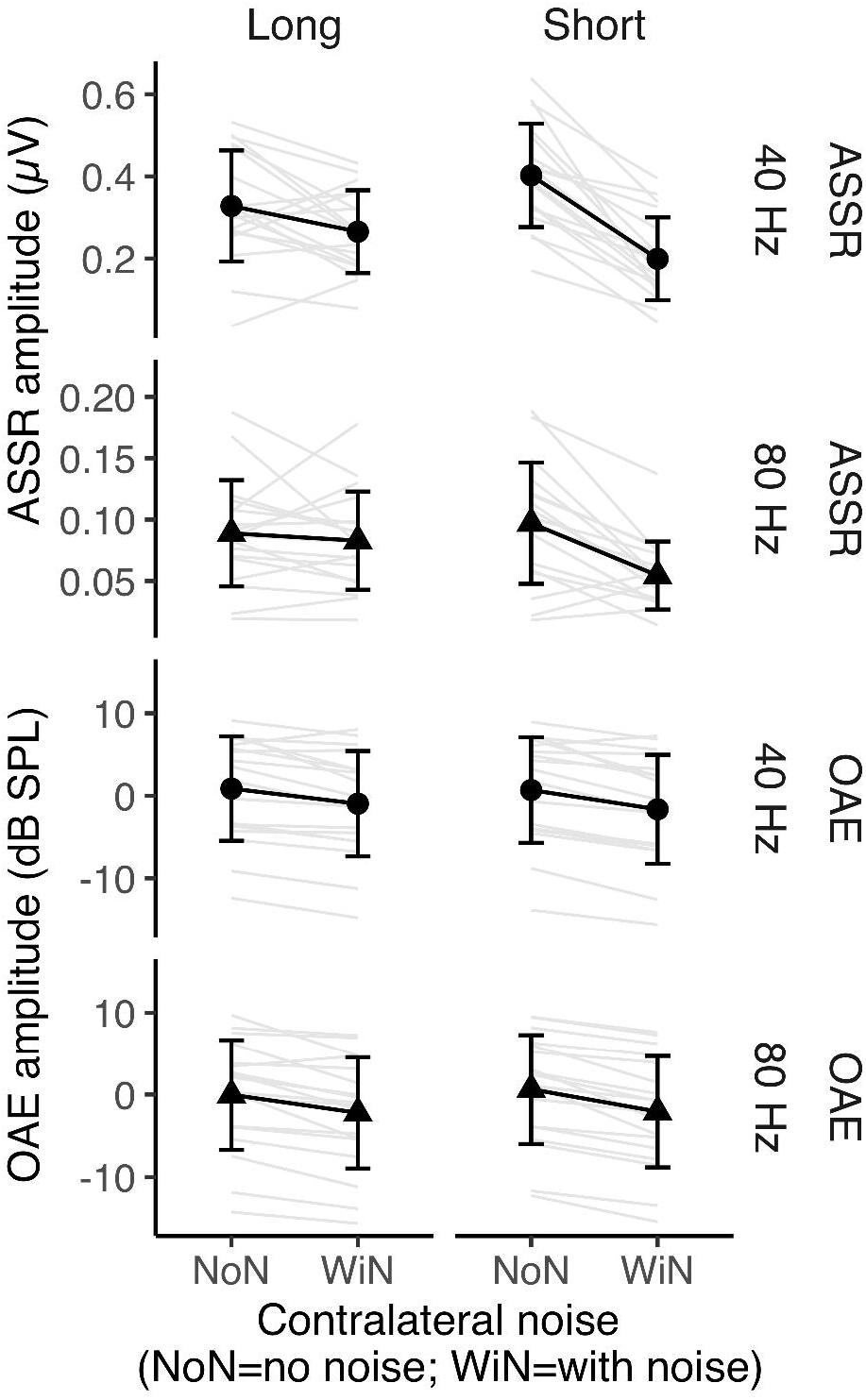
Response amplitude change with contralateral noise. ASSR in the top four panels and OAEs in the bottom four panels. Columns separate long and short duration conditions and rows separate 40 and 80 Hz click-rates. Black circles (40 Hz click-rate) and black triangles (80 Hz click-rate) indicate group means and grey lines represent individual participants. Error bars represent ± one standard deviation. Asterisks denote a significant difference in amplitude between with- (WiN) and no-noise (NoN) conditions.

A 3-way repeated measures ANOVA on OAE amplitude showed no significant 3-way interaction but the two-way interaction between duration and click rate (*F*[1, 17]= 13.2, *p*=0.002) and the two-way interaction between duration and noise (*F*[1, 17]=8.81, *p*=0.008) were significant. Further, OAE amplitude also varied as a main effect of noise (*F*[1, 17] = 58.5, *p*<0.001) suggesting and peripheral inhibition through MOCR activation, as expected. Post-hoc *t*-tests suggest the effect of noise on OAE amplitude was significant for both short (*p*<0.001) and long (*p*<0.001) durations when averaged over both rates.

To investigate the influence of MOCR inhibition at the periphery (OAEs) on the inhibition at brainstem (80 Hz ASSR) and cortical (40 Hz ASSR) levels, we performed correlations. These relations are plotted in Fig. 4. ASSR and OAE inhibition were not correlated for the 40 Hz short condition (*p*=0.454), the 40 Hz long condition (*p*=0.962), and the 80 Hz long condition (*p*=0.981). However, the change in OAEs and ASSRs was positively correlated in the 80 Hz short duration condition (*p*=0.045). Although the *p*-value is only marginally significant for this correlation, the coefficient (*r*=0.48) is consistent with a moderate-to-large effect (Cohen, 1988) and the trend in the data (Fig. 4) is quite apparent unlike the other non-significant correlations. This result likely suggests that changes observed in ASSRs generated predominantly at the cortex (40 Hz) are likely not influenced by MOCR-induced OAE changes at the periphery. However, for ASSRs generated predominantly at the brainstem (80 Hz), the changes at the periphery likely influence neural activity when viewed in shorter time intervals, but this washes out when observed over a longer time window.

**Figure 4:**
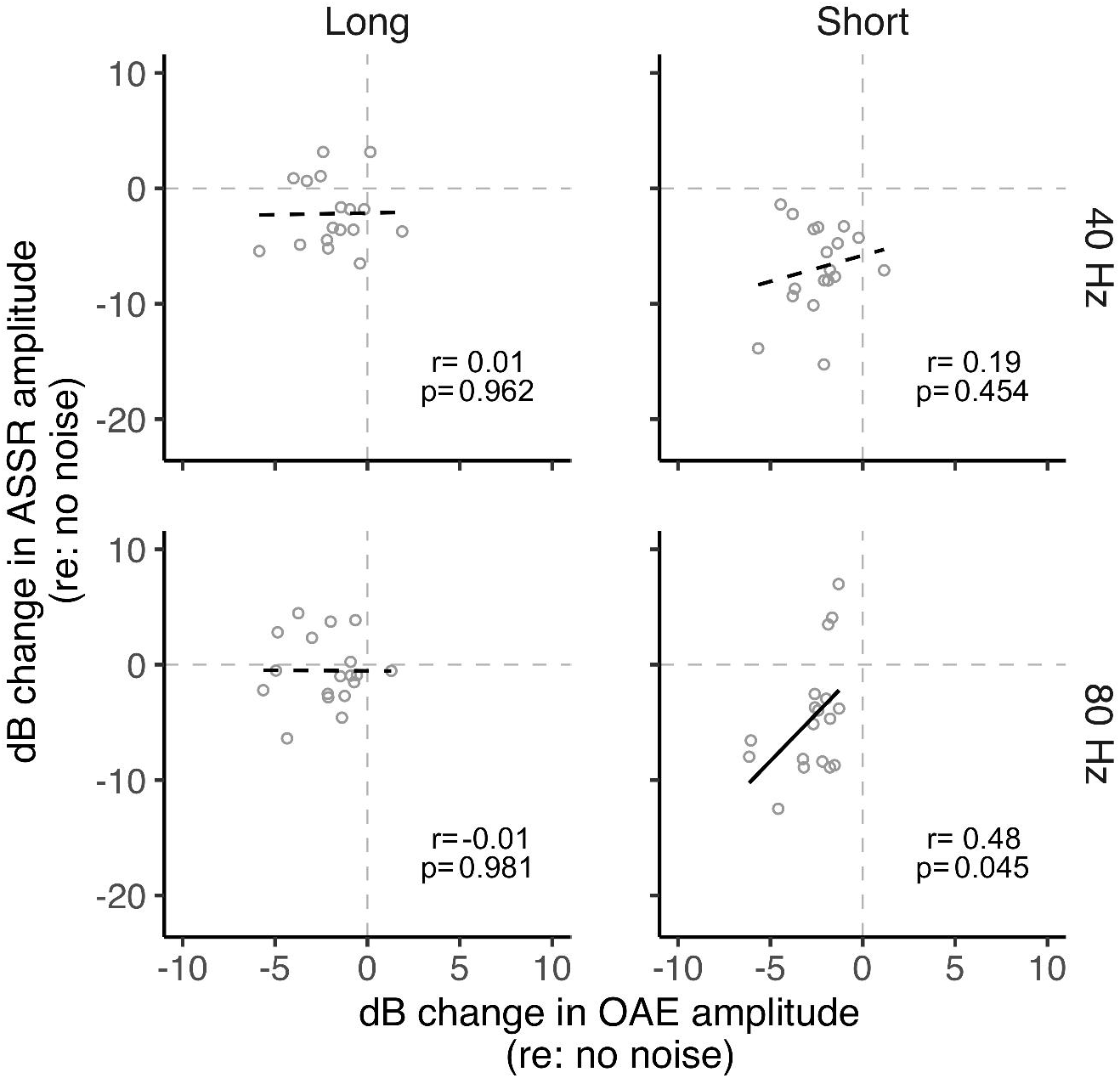
ASSRs vs. OAE amplitude change. (A) 40 Hz click-rate, long stimulus duration (B) 40 Hz click-rate, short stimulus duration (C) 80 Hz click-rate, long stimulus duration (D) 80 Hz click-rate, short stimulus duration. Open circles represent individual participants. A black solid fit line represents a significant relationship between variables. A black dashed fit line represents a nonsignificant relationship between variables.

The relation between the short and long durations for ASSRs and OAEs in the magnitude of inhibition are shown in Fig. 5. There was a significant positive correlation between OAE inhibition in short and long durations observed for both 40 Hz (*p*<0.001) and 80 Hz (*p*=0.001) rates. This indicates that individuals exhibited similar OAE inhibition magnitudes in long and short durations, likely due to a single mechanism, the MOCR, influencing activity at the periphery. However, this relationship was not observed for the ASSR inhibition between short and long for 40 Hz (*p*=0.770) or 80 Hz (*p*=0.346), suggesting differential effects of the eliciting contralateral noise depending on the duration of stimulus presentation.

**Figure 5:**
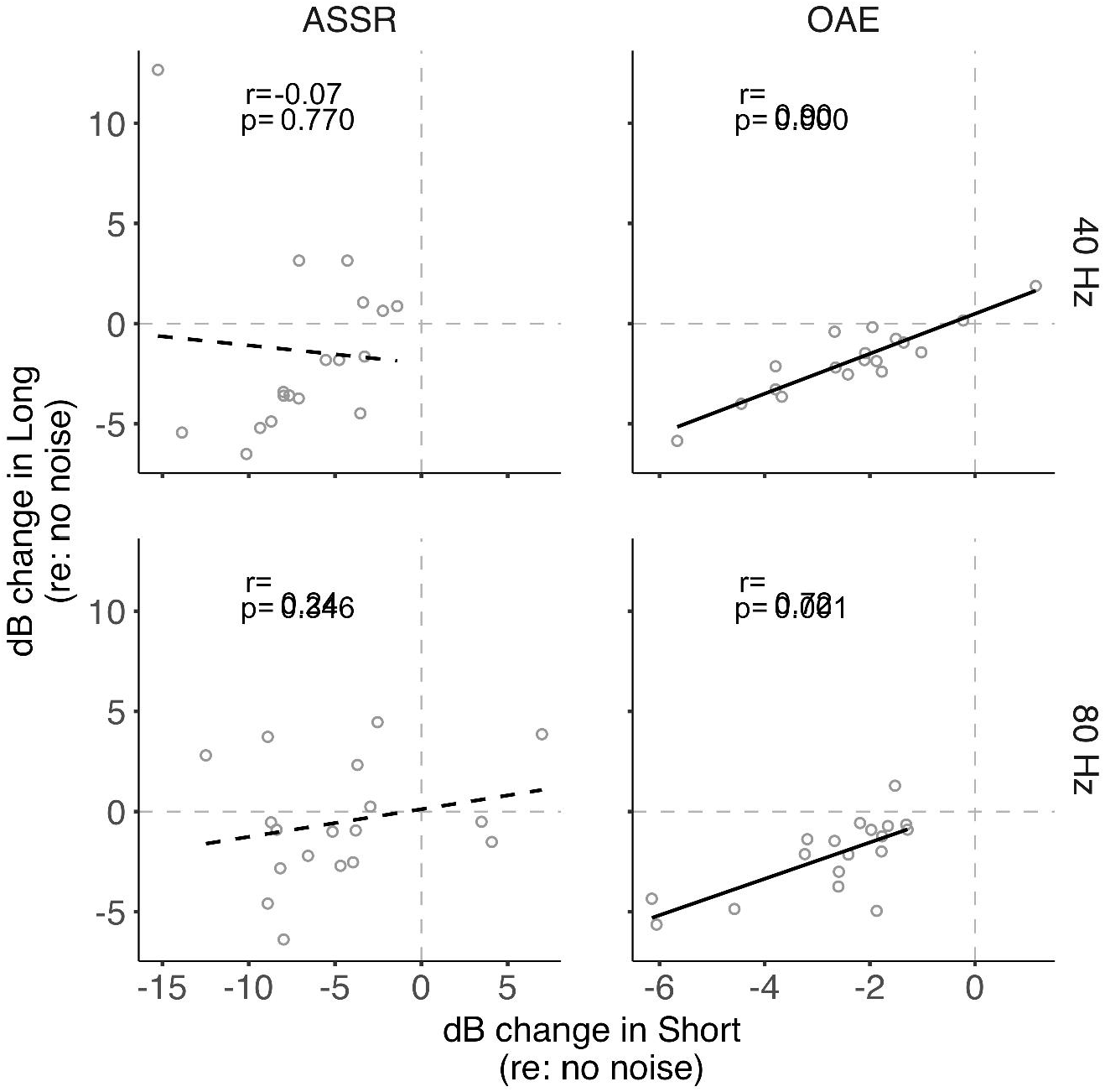
Long vs. short duration. Amplitude changes in the long stimulus durations as a function of amplitude change in the short stimulus durations are plotted for (A) 40 Hz click-rate, ASSRs (B) 40 Hz click-rate, OAEs (C) 80 Hz click-rate, ASSRs (D) 80 Hz click-rate, OAEs. Open circles represent individual participants. A black solid regression line indicates a significant relationship between the two variables. A black dashed regression line indicates a non-significant relationship between the two variables.

We also compared inhibition between the two click rates within short and long durations and separately for ASSRs and OAEs to investigate if any relationships exist between rates. That is to test if responses generated at various levels of the auditory pathway are related in the manner they are measured in this study. The comparisons and correlations are plotted in Fig. 6. For ASSRs, there was no correlation between the inhibition at both click rates for ASSR short (*p*=0.299) and long (*p*=0.434) durations, as expected. This likely indicates that amplitude changes observed in cortical-dominated (40 Hz) ASSRs are independent of brainstem-dominated (80 Hz) ASSR changes in both time durations. For OAEs, while there was a significant correlation between 40 and 80 Hz in the short condition (*p*=0.020), the correlation in the long condition was not significant (*p*=0.961), consistent with the interaction between rate and duration in the ANOVA for OAEs. Taken together, these results align with our hypothesis that brainstem mechanisms likely compensate for self-imposed peripheral inhibition.

**Figure 6:**
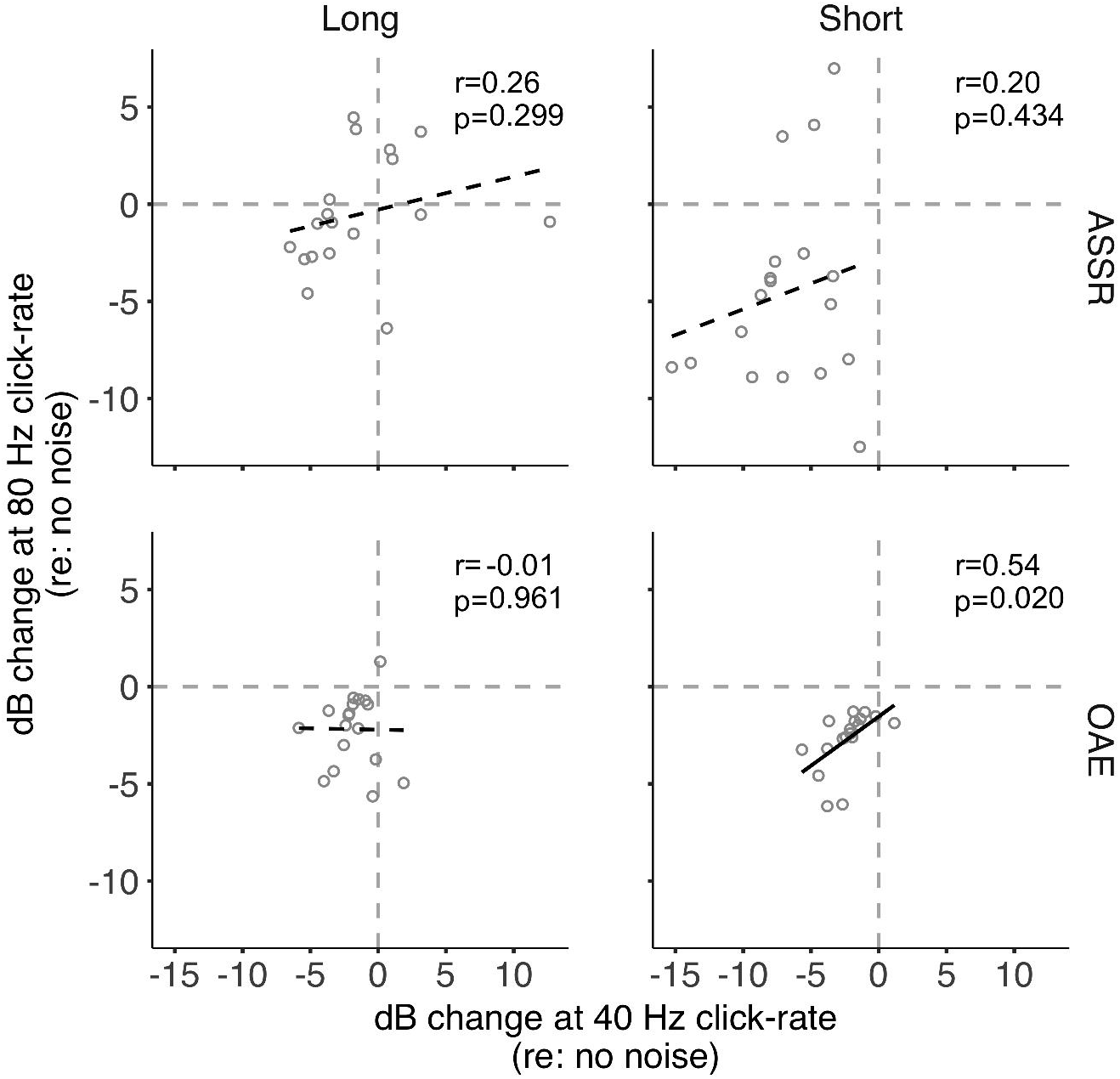
40 Hz vs. 80 Hz. Amplitude changes at 80 Hz click-rate as a function of amplitude change at 40 Hz click-rate for (A) ASSRs, long stimulus duration (B) ASSRs, short stimulus duration (C) OAEs, long stimulus duration (D) ASSRs, short stimulus duration. Open circles represent individual participants. A black solid regression line indicates a significant relationship between the two variables. A black dashed regression line indicates a non-significant relationship between the two variables.

## Discussion

By monitoring cochlear, brainstem and cortical activity with and without MOCR activation in short and long stimulus durations (1.5 s and 4 mins), we found (1) significant inhibition at the periphery and cortex, but not at the brainstem, with long duration stimuli, consistent with previous studies (Özdamar & Bohórquez, 2008; Kawase et al., 2012; Maki et al., 2009; Ross et al., 2005, 2012; Usubuchi et al., 2014), and (2) significant inhibition at the periphery, cortex and notably the brainstem, with short duration stimuli. Below, we discuss these results based on known gain compensation circuits at the brainstem reported in animal models (Hockley et al., 2021; Fujino and Oertel, 2001) and potential alternative reasons.

### Gain compensation in the brainstem

In contrast to the 40 Hz ASSR, the 80 Hz ASSR demonstrates differential inhibition based on the stimulus duration. Methodologically, the total duration of stimulus presentation was equalized between the long and short duration conditions and therefore does not warrant the observed differential effect. In addition, at the periphery, the MOCR inhibition of OAEs across rates and durations is indistinguishable. Given the lack of methodological differences and similarity in peripheral MOCR inhibition between the two durations, the differential effect of noise on the 80 Hz, but not 40 Hz ASSR, can be conjectured to arise either (1) from the difference in the sensitivity of the respective generators to contralateral noise or (2) second-degree factors that influence the generated ASSR differentially between brainstem and more rostral centers.

Previous studies have demonstrated generator-specific effects of contralateral noise. Notably, Özdamar & Bohórquez (2008) showed robust attenuation of later occurring components (Pb) of the middle latency responses elicited at 40 Hz, presumably generated in the cortex. However, such an effect was not observed for concurrently measured ABR wave-V, generated in the brainstem. Similarly, Galambos and Makeig (1992) reported significant attenuation of click-evoked 40 Hz ASSR but not concurrently recorded ABRs. Maki et al. (2009), like the present study compared attenuation of 40 and 80 Hz ASSRs to contralateral noise. They reported reduction of 40 Hz ASSRs but no effect of the contralateral noise on 80 Hz ASSRs. Speculations for such disparity in noise effects on responses generated at the brainstem vs. the cortex has been attributed to one of two reasons. It is possible that noise effects must occur more rostral to the ABR generation site (brainstem; Galambos and Makeig, 1992; Usubuchi et al., 2014), i.e., the generators themselves must be different or that any peripheral inhibition by the MOCR is insignificant for neural measures (Özdamar & Bohórquez, 2008).

The speculation that noise-mediated inhibition of neural responses, also referred to as central masking, occurring only rostral to the brainstem (Galambos and Makeig, 1992; Usubuchi et al., 2014) does not explain the differential noise effects for different types of cortical responses. For instance, although Usubuchi et al. (2014) found significant inhibition for both 20 and 40 Hz neuromagnetic ASSRs, the effect of contralateral noise was more pronounced for the 40 Hz response. While comparing the neuromagnetic 40 Hz ASSRs and the sensory-driven N100 response, Kawase et al. (2012) found robust inhibition of 40 Hz with contralateral noise but no effect on the N100. However, Okamoto et al. (2005) was demonstrated inhibition of the same neuromagnetic N100 like many EEG estimates of N100 (e.g., Salo et al., 2003; Rao et al., 2020). A notable difference between Okamoto et al. (2005) and Kawase et al. (2012) is that Okamoto et al. (2005), apart from studying notch width effects of the noise, use a paradigm that is more akin to our short interval condition where the effect of noise was captured on the test stimulus within 0.5s. In general, there appears to be variable effects of contralateral noise even within responses generated in the cortex and thalamocortical regions. This suggests that other second-degree factors, likely both stimuli-based (e.g., stimulus duration) and physiology-based (e.g., phase reset hypothesis on 40 Hz generation; Ross et al., 2005), are at play in the contralateral noise-mediated inhibitory effects on cortical activity. These factors might also apply for the differences in noise effects observed between cortex-vs. brainstem-generated ASSRs in the present study simply by virtue of the two generators being different. The different generator, however, does not support contralateral noise sensitivity to structures only rostral to the brainstem.

The argument for OHC inhibition by the MOCR being insignificant for neural inhibition (Özdamar & Bohórquez, 2008) also does not explain the disparity in noise effects across neural response generators because (1) the present study results show significant attenuation of 80 Hz ASSRs when acquired in short time intervals and (2) attenuation of auditory nerve compound action potentials is an established marker of MOCR activity on the afferent auditory pathway (Galambos, 1956; Guinan and Gifford, 1988; Liberman, 1989). As such, MOCR-mediated inhibition of OHC amplifier gain is indeed reflected in the auditory nerve response. Therefore, it is still perplexing that the MOCR-mediated attenuation is absent for responses generated at the brainstem when averaged over minutes, especially when it is present (1) at areas rostral to the brainstem, and (2) when responses at the brainstem are considered in shorter time intervals.

Our results may, for the first time, provide a reasoning for the perplexing “insignificant” effect of the MOCR on brainstem generated evoked potentials. Consistent with our hypothesis, we posit that the MOCR-mediated peripheral inhibition probably does become insignificant at the brainstem because mechanisms in the brainstem compensate for the loss in input at the periphery. Known feedback circuitry in the cochlear nucleus support our hypothesis. Based on positive feedback loops between the cochlear nucleus and the superior olivary complex identified in previous animal studies, the likely candidates for such compensation would be the T-stellate (Fujino & Oertel, 2001; Brown, 2011) and the small cells (Hockley et al., 2021).

Anatomically, both T-stellate and small cells of the CN receive direct inputs from the auditory nerve (Liberman, 1991; Blackburn and Sachs, 1989), project to MOC neurons (Romero and Trussell, 2021; de Venecia, et al., 2005; Benson and Brown, 2006; Thompson and Thompson, 1991; Darrow et al., 2011), and receive collaterals from MOC neurons (Benson and Brown, 1990; 1992; Benson et al., 1995; Fujino and Oertel, 2001). These projections and inputs create positive feedback loops for both cell types (i.e., T-stellate-MOC-T-stellate and small cell-MOC-small cell). However, Hockley et al. (2021) reported that activating MOC neurons increased excitation of small cells, but not T-stellate cells. This is intriguing given prior evidence for profuse amounts of MOC collaterals to multipolar/chopper/T-stellate cells (Fujino & Oertel, 2001; Benson and Brown, 1990; review: Brown, 2011). Hockley et al. (2021) do not report how activities of the two CN cell types vary across time scales (e.g., seconds to minutes). Regardless, it is clear that MOC neurons retroactively excite CN neurons, quite likely the same neurons that excite them. In addition, both the MOC neurons and T-stellate cells project to the inferior colliculus (IC) (Schofield, 2001; Okoyama et al., 2006), and a recent study suggests that inputs from CN and IC together optimize MOC activity (Romero and Trussell, 2021). The projection to the IC is important because it is a major generation site for 80 Hz ASSRs (Bidelman, 2018; Herdman et al., 2002; Kuwada et al., 2002; review: Dimitrijevic and Ross, 2008) and is in the vicinity of the ABR wave-V generator (Moller and Jannetta, 1982).

### Relevance of gain compensation at the brainstem

The putative positive MOCR feedback loops for both cell types, the T-stellate and small cells, are thought to act as ‘efferent copies’ of MOCR inhibition at the periphery and likely compensate for the reduced input (Brown, 2011; Fujino and Oertel, 2001). This gain compensation could be critical for at least three reasons. First, a reduction of input at the periphery will decrease excitation of the MOCR and the MEMR, which will in turn limit their functional ability to protect vulnerable cochlear hair cells from acoustic overexposure (Rajan, 1988; Liberman, 1990; Lauer and May, 2011; Brown, 2011; Fujino and Oertel, 2001). Second, a reduction of input at the periphery might negatively impact central gain, and possibly the tuning of the T-stellate cells (Rhode and Greenberg, 1994) as it would alter the input to D-stellate cells. By providing inhibitory input to the T-stellate cells, the D-stellates provide a balance in gain in the CN among their various other functions (Ferragamo et al., 1998; Oertel et al., 2011; Rhode and Greenberg, 1994). Reduced T-stellate cell inhibition, combined with the recently discovered excitatory loop gain within T-stellate cells (Cao et al., 2019) could lead to tinnitus and hyperacusis when left unchecked (Hockley et al., 2021). Third, compensating for reduced peripheral input likely restores, and possibly enhances, the fidelity of the sound level, specifically the spectral peaks as encoded by T-stellate cells (Blackburn and Sachs, 1990; May et al., 1998; Oertel et al., 2011) and small cells (Hockley et al., 2021). At a population level, both T-stellate cells and small cells encode spectral peaks and have been identified to be critical for speech perception (Hockley et al., 2021; Oertel et al., 2011; May, LePrell, and Sachs, 1998; Blackburn and Sachs, 1990). The MOC neurons only provide cholinergic input to the T-stellate, not D-stellate cell, which is thought to lead to selective enhancement of spectral peaks and not valleys, improving the overall SNR in the system (Fujino and Oertel, 2001). Further, T-stellate and small cells respond optimally at moderate to high stimulus levels, like the elicitor used in the present study (60 dB SPL; Lai, Winslow, and Sachs, 1994; Liberman, 1991; Ryugo, 2008) and are, therefore, likely to be reflected in our experimental approach.

Taken together, the compensation mechanisms likely reflected in our results are critical for maintaining homeostasis, continued protection of peripheral structures, and prevent peripheral inhibition from degrading the encoding of important acoustic information. By contrasting short vs. long duration conditions, our results provide a potential non-invasive window into these mechanisms. As with any non-invasive markers, their true physiological origins must be established using direct and likely invasive studies of these systems. Specifically, future studies capable for selectively silencing the feedback from the T-stellate and small cells in the CN to the MOCR may provide the most conclusive evidence for our hypothesized gain compensation mechanism.

### If the brainstem compensates, why is inhibition still present at the brainstem and the cortex?

If the inhibition of peripheral input is compensated for in the CN, the inhibition of brainstem-generated (80 Hz) ASSRs observed in the short duration condition reveals a time course to this compensation. Currently, there are no studies that directly describe the physiological time course of T-stellate/small cell-MOCR-mediated gain compensation. Nevertheless, the pattern of results in this study may be explained by the established kinetics of the MOCR pathway. It can be conjectured that inhibition of auditory nerve inputs by the MOCR causes an initial reduction in inputs to the CN, likely on a scale of several tens to a few hundred milliseconds, commensurate with the MOCR activation time course of 200-250 ms and the roughly 400 ms it to reach steady state (Backus and Guinan, 2006; Boothalingam et al., 2021). Considering that both the MOCR and T-stellate cells integrate energy over time and continue to be excited even after stimulus cessation, the gain compensation could happen over a few seconds. This initial reduction in peripheral input is reflected as reduced brainstem-dominated ASSR amplitude (80 Hz) in the short duration condition and is somewhat supported by the positive correlation with MOCR inhibition of OAEs in this condition (Fig 4D).

When contralateral noise was introduced, a reduction in cortex-generated (40 Hz) ASSR amplitude was observed with both long stimulus durations consistent with previous studies (Galambos & Makeig, 1992; Ross & Fujioka, 2016; Maki et al., 2014; Usubuchi et al., 2009), as well as with short stimulus durations. However, the lack of significant correlation between the change in ASSR amplitude for 40 Hz and OAEs (Fig 4) suggest that the reduction in 40 Hz ASSR amplitude is unlikely to be related to the MOCR-mediated peripheral inhibition. This finding corroborates the findings of Mertes and Leek (2016) and Mertes and Potocki (2022) – significant reduction in 40 Hz ASSR without any correlation with the inhibition of OAEs. The reduction of 40 Hz ASSR amplitude may instead be explained by an interruption in thalamocortical loop resonance induced by contralateral noise. Desynchronization of 40 Hz ASSR, associated with a temporary decrease in the amplitude of oscillatory signal power in response to a concurrent stimulus, was similarly observed by Ross and colleagues (2005; 2012). ASSR desynchronization is a general reaction to both new and changing stimuli (Ross et al., 2005) thought to act as a reset to the adaptation of auditory processing (Rogers & Bregman, 1998). The reduction in 40 Hz ASSR amplitude may reflect this temporary desynchronization, which is more evident when averaging amplitude over short compared to long durations.

### Clinical implications of a non-invasive window into brainstem gain compensation

Considering the role that peripheral input to the CN plays in the gain compensation mechanism, the impact of hearing loss should be further investigated for diagnostic/therapeutic implications and for uncovering the mechanism’s ecological purpose. For instance, in the case of sensorineural hearing loss resulting from outer hair cell damage, MOCR inhibition would not affect the outer hair cell activity, but the gain compensation may still enhance T-stellate and small cell excitation. This process may result in a net positive excitation – overcompensation – in the CN. This overcompensation may contribute to auditory disorders characterized by hyperactivity such as tinnitus and hyperacusis (Noreña, 2011; Zeng, 2013; Cao et al., 2019; Hockley et al., 2021).

MOCR function is important to consider in a clinical setting as it has been hypothesized to improve speech perception in noise (Giraud et. al., 1997: Kumar and Vanaja, 2004; Mishra and Lutman, 2014). Prior arguments for an MOCR role in speech perception were based only on its ability to restore the dynamic range of the auditory nerve (Guinan, 2006). Evidence of MOCR collateral activity in CN – enhancing T-stellate and small cell output – further strengthens the role of MOCR in speech perception (Fujino and Oertel, 2001; Hockley et al., 2021). OAEs are typically used to measure the MOCR strength in normal hearing individuals. Given that OAEs rely on OHC activity, hearing loss due to OHC damage emphasizes the need for alternative measures of MOCR function. While ASSRs appear to be a promising alternative, our findings indicate that the time scale and generation site at which inhibitory effects are compensated for must be carefully considered. For instance, the contralateral noise-mediated inhibition of 40 Hz ASSRs (Mertes and Leek, 2016; Mertes and Potocki, 2022; Usubuchi et al., 2014; Maki et al., 2009) may not reflect MOCR inhibition of OHC activity (Fig 4) as any reduction in peripheral input to the cortex is likely iteratively compensated for at the brainstem.

In summary, our findings offer a potential new window into a gain compensation mechanism in the brainstem previously identified in animal models. Additionally, this study demonstrates the ability of the 80 Hz ASSR to measure MOCR function using short duration stimuli. Our methods and corresponding results also emphasize the importance of timescale consideration for future research utilizing ASSRs to measure the MOCR.

## Acknowledgments

This work was supported by the Office of the Vice Chancellor for Research and Graduate Education, University of Wisconsin-Madison. Portions of this work were presented at the 2019 American Auditory Society Meeting, Scottsdale, AZ.

